# iCodon: ideal codon design for customized gene expression

**DOI:** 10.1101/2021.05.06.442969

**Authors:** Santiago Gerardo Medina-Muñoz, Michay Diez, Luciana Andrea Castellano, Gabriel da Silva Pescador, Qiushuang Wu, Ariel Alejandro Bazzini

## Abstract

Messenger RNA (mRNA) stability substantially impacts steady-state gene expression levels in a cell. mRNA stability, in turn, is strongly affected by codon composition in a translation-dependent manner across species, through a mechanism termed codon optimality. We have developed iCodon (www.iCodon.org), an algorithm for customizing mRNA expression through the introduction of synonymous codon substitutions into the coding sequence. iCodon is optimized for four vertebrate transcriptomes: mouse, human, frog, and fish. Users can predict the mRNA stability of any coding sequence based on its codon composition and subsequently generate more stable (optimized) or unstable (deoptimized) variants encoding for the same protein. Further, we show that codon optimality predictions correlate with expression levels using fluorescent reporters and endogenous genes in human cells and zebrafish embryos. Therefore, iCodon will benefit basic biological research, as well as a wide range of applications for biotechnology and biomedicine.

## Introduction

The genetic code is degenerate, as most amino acids are encoded by multiple codons (Gouy & Gautier, 1982). The codons encoding for the same amino acid are called synonymous or silent codons. Long regarded as interchangeable, these codons are not equivalent from a regulatory point of view (Gouy & Gautier, 1982). Synonymous codons are used with different frequencies in the coding genome, a phenomenon known as codon usage bias (Sharp & Li, 1987). Moreover, synonymous codon substitutions can dramatically affect messenger RNA (mRNA) stability and therefore protein production (Boël et al., 2016; de Freitas Nascimento, Kelly, Sunter, & Carrington, 2018; Gouy & Gautier, 1982; Q. Wu et al., 2019). Recent studies have revealed that translation strongly affects mRNA stability *in cis* in a codon-dependent manner in vertebrates (Bazzini et al., 2016; Mishima & Tomari, 2016; Q. Wu et al., 2019) as well as in other species (Boël et al., 2016; Burow et al., 2018; de Freitas Nascimento et al., 2018; Harigaya & Parker, 2016; Jeacock, Faria, & Horn, 2018; Presnyak et al., 2015; Radhakrishnan et al., 2016), a process referred as codon optimality (Presnyak et al., 2015). Codon optimality is the most pervasive mechanism underlying mRNA stability in yeast (Cheng, Maier, Avsec, Rus, & Gagneur, 2017) and vertebrates (Medina-Muñoz et al., 2021). Specifically, to determine the regulatory strength of codon optimality in vertebrates, we have recently developed a machine learning model that predicts mRNA stability based on codon composition (Medina-Muñoz et al., 2021). Trained with multiple profiles of mRNA stability for thousands of genes obtained from human (Q. Wu et al., 2019) and mouse cells (Herzog et al., 2017), as well as *Xenopus* and zebrafish embryos (Bazzini et al., 2016; Medina-Muñoz et al., 2021), this model has revealed that codon composition is a major determinant of mRNA stability during early embryogenesis and dictates mRNA levels in conjunction with other *cis*-regulatory elements (e.g., microRNA and m^6^A) in human and mouse cells as well as in zebrafish and *Xenopus* embryos (Medina-Muñoz et al., 2021). Therefore, we hypothesized that the model could be used as a tool for the design of synonymous coding sequences with differing stability characteristics depending on the desired application.

Existing methods to perform codon optimization are mainly based on codon usage bias (Burgess-Brown et al., 2008; Fuglsang, 2003; Puigbo, Guzman, Romeu, & Garcia-Vallve, 2007; G. Wu, Bashir-Bello, & Freeland, 2006). Yet, in vertebrates, weak positive correlations have been observed between codon usage bias and codon optimality (Bazzini et al., 2016; Q. Wu et al., 2019). For example, the codon usage bias for some amino acids (e.g., Arginine and Threonine) differs drastically from codon optimality (Q. Wu et al., 2019). Therefore, a method for codon optimization using codon optimality represents a novel approach for *in silico* gene design.

Here, we developed a tool named iCodon (www.iCodon.org) that optimizes coding regions with synonymous codon substitutions to increase mRNA stability and therefore protein expression (e.g., to design highly expressed reporters), or deoptimize sequences with synonymous codon substitutions to decrease mRNA stability (e.g., to design a sequence with decreased expression). iCodon uses a predictive model of mRNA stability (Medina-Muñoz et al., 2021) as a guide for supervising the design of sequences. Therefore, iCodon can also be used to visualize the predicted mRNA stability based on the codon composition of any coding sequence.

In summary, iCodon chooses ideal codons for incorporation into designed coding sequences to reach desired gene expression levels. iCodon is available as an R package (https://github.com/santiago1234/iCodon) and an interactive web interface www.iCodon.org (https://bazzinilab.shinyapps.io/icodon/).

## Results

### iCodon predicts gene expression based on codon composition and designs new variants based on synonymous substitutions

Our machine learning model to predict mRNA stability as a function of codon composition (Medina-Muñoz et al., 2021) was trained with mRNA stability profiles from zebrafish and *Xenopus* embryos (Bazzini et al., 2016; Medina-Muñoz et al., 2021), human cell lines (Q. Wu et al., 2019), and mouse embryonic stem cells (Herzog et al., 2017). We hypothesized that our model could be used to supervise the design of coding sequences with customized host mRNA stabilities based on synonymous codon choice.

First, to test the sensitivity of the predictive model to capture synonymous substitution effects on gene expression, we analyzed previously published reporter sequences designed to generate identical peptides but differing in codon choice (Q. Wu et al., 2019). These reporters encode for mCherry followed by a ribosome skipping sequence (P2A) (de Felipe et al., 2006; Donnelly et al., 2001) and a region enriched in optimal (stabilizing) or non-optimal (destabilizing) synonymous codons (**Figure 1A**). Importantly, and due to the P2A sequence, mCherry production is independent of potential protein folding differences that may arise for the peptides encoded by the variable region (optimal or non-optimal). Previously, we have shown that mRNA levels and fluorescence intensities in transfected human 293T cells correlate with the proportion of optimal codons in the reporter sequences (Q. Wu et al., 2019). Here, we found that the model correctly estimated the expression profile of these reporters in transfected human 293T cells (p value < 2.2×10^−16^, Pearson correlation test) (**Figure 1B**). In contrast, a reporter optimized according to codon usage parameters (IDT Codon Optimization tool, www.idtdna.com) displayed reduced fluorescence intensity and predicted stability when compared to reporters enriched in optimal codons **(Figure 1B)** (Synonymous_3 or Synonymous _4 vs Synonymous_Usage, p < 1.0×10^−07^, paired t-test). Therefore, our model is able to predict the impact of synonymous codon changes on gene expression.

**Figure 1.**
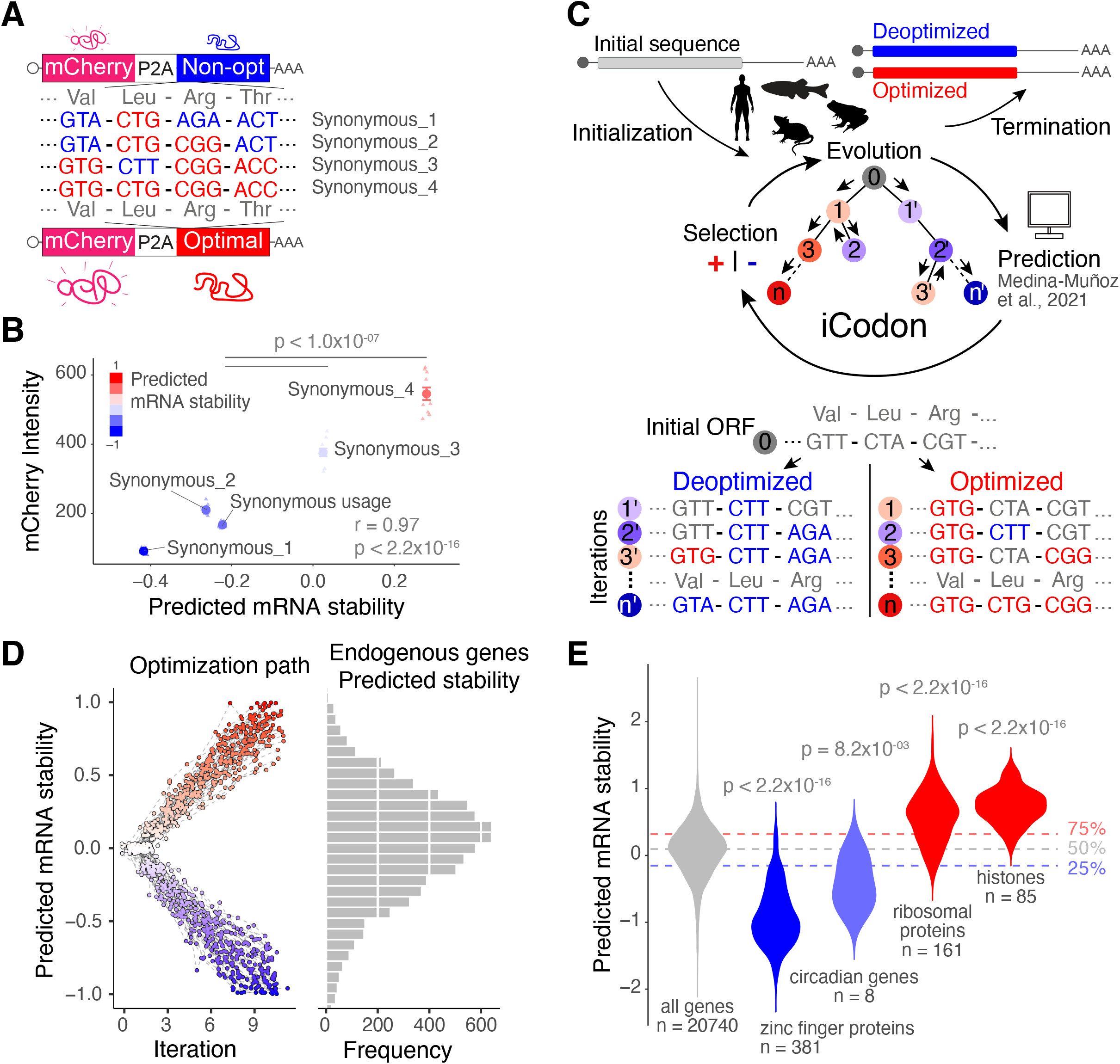
iCodon predicts gene expression based on the codon composition and designs new variants based on synonymous mutations. **(A)** Diagram of the synonymous reporters, differing only in synonymous mutations with different regulatory effects on mRNA stability. Each mRNA contains the coding sequence of mCherry fluorescent protein followed by a ribosome-skipping sequence (P2A) and a coding region that differs in the proportion of optimal and non-optimal codons but encodes the same peptide (synonymous mutations). **(B)** Scatter plot showing that fluorescence intensity of synonymous reporters in 293T transfected cells correlates with predicted mRNA stability (r = 0.97, p < 2.2×10−16, Pearson correlation test). **(C)** Diagram depicting the algorithm for codon optimization, iCodon: An initial coding sequence is provided by the user. Evolution: the algorithm generates variant sequences by introducing random synonymous mutations. Prediction: the machine learning model predicts the mRNA stability of each variant based on the codon composition. Selection: the algorithm selects the sequences with the highest or lowest mRNA stability depending on the direction of optimization. Iteration: this process is repeated multiple times producing an optimization path that generates a gradient in mRNA stability level. **(D)** A random group of 50 human genes with predicted intermediate mRNA stability was selected and optimized and deoptimized by iCodon. The x-axis is the iteration number, and the y-axis is the predicted mRNA stability. The circles connected by a dash-line show the optimization or deoptimization path for each gene. The histogram on the right is the mRNA stability distribution for endogenous human genes. **(E)** Violin plot showing predictions of mRNA stability of selected groups of genes compared to all genes in the human transcriptome. The horizontal lines show the lower, middle and upper quartiles of the predicted mRNA stability of all genes. P values and number of genes (n) are indicated.

Next, we created iCodon by coupling the predictive model (Medina-Muñoz et al., 2021) to an evolutionary algorithm (**Figure 1C**), whereby a given sequence accumulates synonymous mutations through multiple selective iterations (see Methods section). The predictive model is used to supervise the selection of new variants with increased (optimized) or decreased (deoptimized) gene expression (**Figure 1C**).

We ran a simulation to optimize and deoptimize a group of 50 randomly chosen human genes with intermediate mRNA stability (**Figure 1D**). This simulation revealed three important aspects. First, these genes can be optimized or de-optimized to reach the predicted mRNA stabilities of the most or least stable genes in the transcriptome, respectively (**Figure 1D**). Second, the algorithm generates a range of intermediate sequences that fall along a gradient of predicted mRNA stabilities (**Figure 1D**). And third, the degree of optimization achieved for most genes is predicted to affect mRNA abundance by orders of magnitude ranging from 10- to 100-fold (**Supplementary Figure 1A**) (Ross, 1995). These results highlight the potential of iCodon to design coding sequences that display wide stability profiles through synonymous codon substitutions.

### iCodon as an mRNA stability predictor tool

Knowing the predicted stability of particular genes based on the coding sequence can be an entry point to understand its function or regulation mode. Pertinent to this, we have generated transcriptome wide predictions of mRNA stability for endogenous genes in human, zebrafish, *Xenopus*, and mouse (**Supplemental Tables 1-4**). For example, Gene Ontology term analysis of the top 100 most optimal human genes showed an enrichment of pathways related to translation. Interestingly, the same analysis of the top 100 most non-optimal genes revealed an enrichment of transcription factors (**Supplemental Table 5**). Moreover, transcription factors belonging to zinc finger proteins (n=381) as well as core circadian genes (n=8) (Takahashi, 2017) displayed a significant lower stability score compared to the stability of the transcriptome (p < 2×10^−16^ and p < 8.2×10^−03^, respectively, unpaired t-test) (**Figure 1E**). Contrary, histones (n=85) and ribosomal proteins (n=161) showed a significant higher stability score compared to the stability of the transcriptome (p < 2×10^−16^ and p < 2×10^−16^, respectively, unpaired t-test) (**Figure 1E**). Therefore, addressing the predicted stability of a given mRNA could be useful to hypothesize about gene function and evolution. For instance, it can be proposed that core circadian genes might have been under evolutionary pressure to be unstable in order to have oscillatory expression. Therefore, researchers may use iCodon to dissect the stability of their desired gene based on its codon composition.

### iCodon generates fluorescent variants with desired expression levels

Next, we tested the potential of iCodon to produce sequences with different expression levels. Using iCodon, we optimized and deoptimized EGFP (enhanced Green Fluorescent Protein) and generated twelve GFP variants ranging in different levels of codon optimality (**Figure 2A**). Interestingly, the predicted mRNA stability correlated with protein fluorescence intensities observed in transfected 293T human cells (r = 0.89, p value < 2.2×10^−16^, Pearson correlation test) (**Figure 2B**). Nearly 50-fold differences in intensity were observed between the GFP variants designed by iCodon with the highest and lowest expression levels (**Figure 2B**). The fluorescence intensity for the most optimal GFP variant (i.e., GFP_12) was not as strong as that observed for EGFP (**Figure 2B**). However, this result was not surprising, as EGFP has been extensively optimized for increased fluorescence (Contag, Olomu, Stevenson, & Contag, 1998; T.-T. Yang, Cheng, & Kain, 1996). Yet, this GFP variant (GFP_12) differs from EGFP by 87 nucleotides and 78 codons (**Figure 2C and Supplementary Figure 1B**), without affecting fluorescence intensity drastically (EGFP vs GFP_12 fold-change = 1.24, p value = 2.0×10^−07^, paired t-test). Moreover, from EGFP, 131 nucleotides (121 codons) were changed to create one of our neutral GFPs (GFP_7), and 127 nucleotides (106 codons) mutations generated our dimmest, destabilized GFP (GFP_2) (**Figure 2C and Supplementary Figure 1B**). However, GFP_7 displayed 14-fold higher fluorescence intensity than GFP_2 (**Figure 2B**). These results show that the type of substitution is more important than the number of mutations to affect gene expression. It is important to mention that some GFP variants, such as GFP_3 and GFP_11, did not necessarily follow the expected expression level (**Figure 2B**). For example, GFP_3 did not produce any detectable GFP expression, which illustrates that there are likely other features (e.g., nucleotide sequence, RNA structure, etc.) affecting protein expression, highlighting the nonequivalent nature of synonymous substitutions. However, iCodon was able to design GFP variants with nearly 50-fold differences in expression levels in human cells (**Figure 2B**).

**Figure 2.**
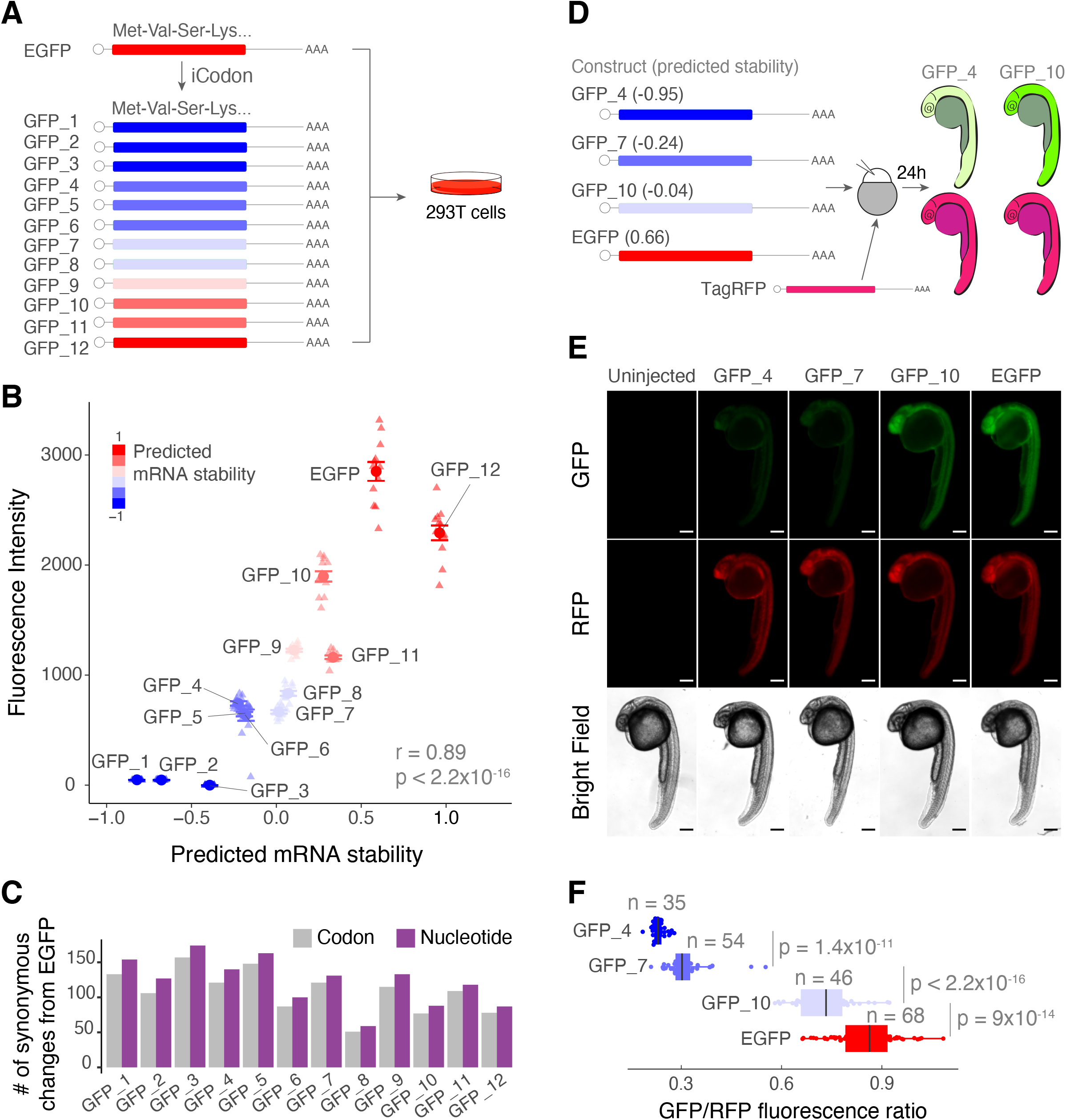
iCodon generates fluorescent variants with desired expression levels. **(A)** Illustration of 12 GFP variants generated by iCodon differing only in synonymous mutations and its predicted mRNA stability. All GFP variants were transfected in 293T cells and the fluorescence was measured by cytometry analysis. **(B)** Scatter plot showing a positive correlation between predicted mRNA stability and GFP fluorescence intensity in 293T transfected cells (r = 0.89, p < 2.2×10^−16^, Pearson correlation test). **(C)** Barplot displaying the number of codon (gray) or nucleotide (purple) changes in all 12 GFP variants compared to EGFP. **(D)** Four GFP variants were co-injected with TagRFP into 1-cell stage zebrafish embryos and imaged 24 hours post fertilization (hpf). The mRNA stability predictions by iCodon in zebrafish are indicated in brackets. **(E)** Microscopy images of injected zebrafish embryos after 24 hpf. Scale bars represent 200 μm. **(F)** Quantification of the differences of fluorescence intensity of GFP relativized by RFP fluorescence from injected zebrafish embryos. P values and replicates (n) are indicated.

Next, to test whether the iCodon predictions correlated with fluorescence expression levels in an *in vivo* model (zebrafish embryos), variants that displayed low (GFP_4), medium (GFP_7) and high (GFP_10) expression in transfected human cells (**Figure 2AB**) were selected. Each variant, as well as EGFP mRNA, were co-injected with TagRFP mRNA as an internal control, into 1-cell stage zebrafish embryos, and GFP/RFP ratio was measured at 24 hours post fertilization (hpf) (**Figure 2D**). The GFP variants displayed profiles of fluorescence intensity (**Figure 2EF**) similar to those observed in human cells **(Figure 2B)**, which correlated with their predicted stability (r = 0.86, p value < 2.2×10^−16^, Pearson correlation test). Together, these results illustrate the ability of iCodon-specified synonymous substitutions to modulate gene expression in both human cells and zebrafish embryos.

### iCodon improves performance of fluorescent AausFP1 variants for expression in vertebrates

We next tested whether iCodon can improve the performance of AausFP1, a recently identified fluorescent protein from *Aequorea. cf. australis* (Lambert et al., 2020). This protein is reported to be 5-fold brighter and more photostable than EGFP (Lambert et al., 2020) but has not yet been optimized for vertebrate expression. Starting with the reported AausFP1 coding sequence (Lambert et al., 2020), four optimized versions were designed by iCodon, as well as two variants designed by a codon usage approach (IDT Codon Optimization Tool, www.idtdna.com) for zebrafish and human expression **(Figure 3A)**. Similar to the GFP results **(Figure 2)**, the stability predicted by iCodon correlated with AausFP1 fluorescence intensities observed in transfected 293T human cells (r = 0.84, p value < 2.2×10^−16^, Pearson correlation test) (**Figure 3B**). Strikingly, a nearly 4-fold change in fluorescence intensity was observed between the original AausFP1 coding sequence and a AausFP1.4 variant optimized by iCodon in 293T cells (p value = 1.6×10^−14^, paired t-test) **(Figure 3B)**. In contrast, only small differences in fluorescence intensity were observed between the original AausFP1 and the variant optimized based on human codon usage (AausFP1vs Human usage, p value = 0.01, paired t-test) in 293T cells **(Figure 3B)**. A variant optimized based on zebrafish codon usage actually displayed reduced fluorescence intensity in transfected human cells (AausFP1vs Zebrafish usage, p value = 1.1×10^−11^, unpaired t-test) **(Figure 3B)**. Similar to the GFP experiment (**Figure 2**), the types of codon substitutions, rather than overall number of substitutions, had the greatest effect on gene expression. For example, the brightest variant optimized by iCodon (AausFP1.4) contained less substitutions than a weaker (lower fluorescence intensity) variant designed according to human codon usage (**Figure 3C and Supplementary Figure 1C**).

**Figure 3.**
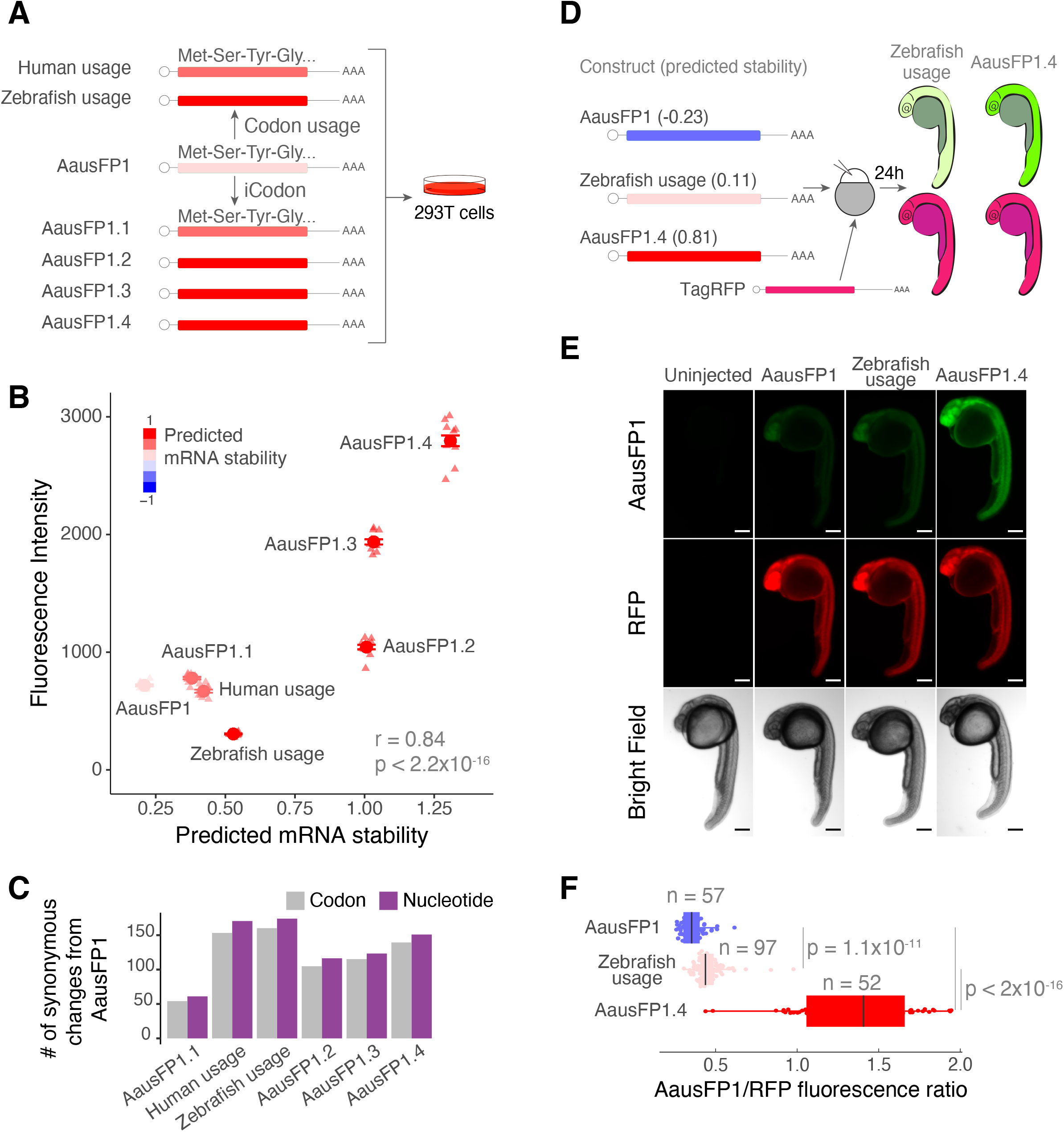
iCodon optimizes fluorescent AausFP1 variants for expression in vertebrates. **(A)** Diagram of the AausFP1 variants optimized by iCodon or by IDT Codon Optimization Tool (codon usage) that were transfected in 293T cells. **(B)** Scatter plot showing a positive correlation between predicted mRNA stability and AausFP1 fluorescence intensity in 293T transfected cells (r = 0.84, p < 2.2×10^−16^, Pearson correlation test). **(C)** Bar plot showing the number of codon (gray) or nucleotide (purple) changes in AausFP1 variants compared to the original sequence. **(D)** Three AausFP1 variants were co-injected with TagRFP into 1-cell stage zebrafish embryos and imaged 24 hours post fertilization (hpf). The mRNA stability predictions by iCodon in zebrafish are indicated in brackets. **(E)** Microscopy images of injected zebrafish embryos after 24 hpf. Scale bars represent 200 μm. **(F)** Quantification of the differences of fluorescence intensity of AausFP1 relativized by RFP fluorescence from injected zebrafish embryos. P values and replicates (n) are indicated.

Next, we tested whether, similar to our GFP experiments **(Figure 2 DEF)**, the iCodon predictions of AausFP1 variants for zebrafish correlate with fluorescence expression levels in zebrafish embryos. Three AausFP1 variants (mRNA) were co-injected with TagRFP mRNA as internal control, into 1-cell stage zebrafish embryos, and AausFP1/RFP ratio was measured at 24 hpf **(Figure 3D)**. We observed a positive correlation between the stability predicted by iCodon and the AausFP1 fluorescence intensities observed in injected zebrafish embryos (r = 0.86, p value < 2.2×10^−16^, Pearson correlation test) **(Figure 3EF)**. Moreover, the optimized variant displayed nearly 4-fold more fluorescence (AausFP1/RFP ratio) than the original AausFP1 (AausFP1.4 vs AausFP1, p value < 2.2×10^−16^, unpaired t-test) and close to 3-fold more fluorescence when compared to the variant optimized according to codon usage (AausFP1.4 vs Zebrafish usage, p value < 2.2×10^−16^, unpaired t-test) **(Figure 3EF)**. In sum, iCodon is able to improve the expression of heterologous coding sequences for use in vertebrate systems.

### Optimized endogenous variant rescues loss-of function phenotypes

The above results demonstrate the ability to modulate the expression of distantly related sequences within a heterologous system. We next hypothesized that iCodon could enhance the expression of coding sequences from within the host genome. Specifically, we asked whether iCodon optimization could improve the ability of endogenous coding sequences to rescue loss-of-function phenotypes that might depend on protein dosage. For this, we first generated a zebrafish line lacking melanin pigmentation (albino phenotype) by targeting gene *slc45a2* with CRISPR/Cas9 (Moreno-Mateos et al., 2015). Then, we generated four *slc45a2* mRNA variants to inject into zebrafish embryos: the original *slc45a2* coding sequence, as well as one variant optimized according to zebrafish codon usage (Codon usage) and two variants designed by iCodon (optimal and non-optimal) for zebrafish expression **(Figure 4A)**. *in vitro* transcribed mRNAs for each *slc45a2* variant were injected into 1-cell stage albino zebrafish embryos. Interestingly, pigmentation was minimally rescued by the unmodified *slc45a2* variant and not rescued by the non-optimal variant after 48 hpf **(Figure 4B)**. However, embryos injected with either optimized variant (according to codon usage or optimality) displayed pigmentation after 48 hpf (**Figure 4B**). These results highlight the ability of iCodon optimization to affect *in vivo* stability of introduced mRNAs, and thereby modulate their therapeutic efficacy.

**Figure 4.**
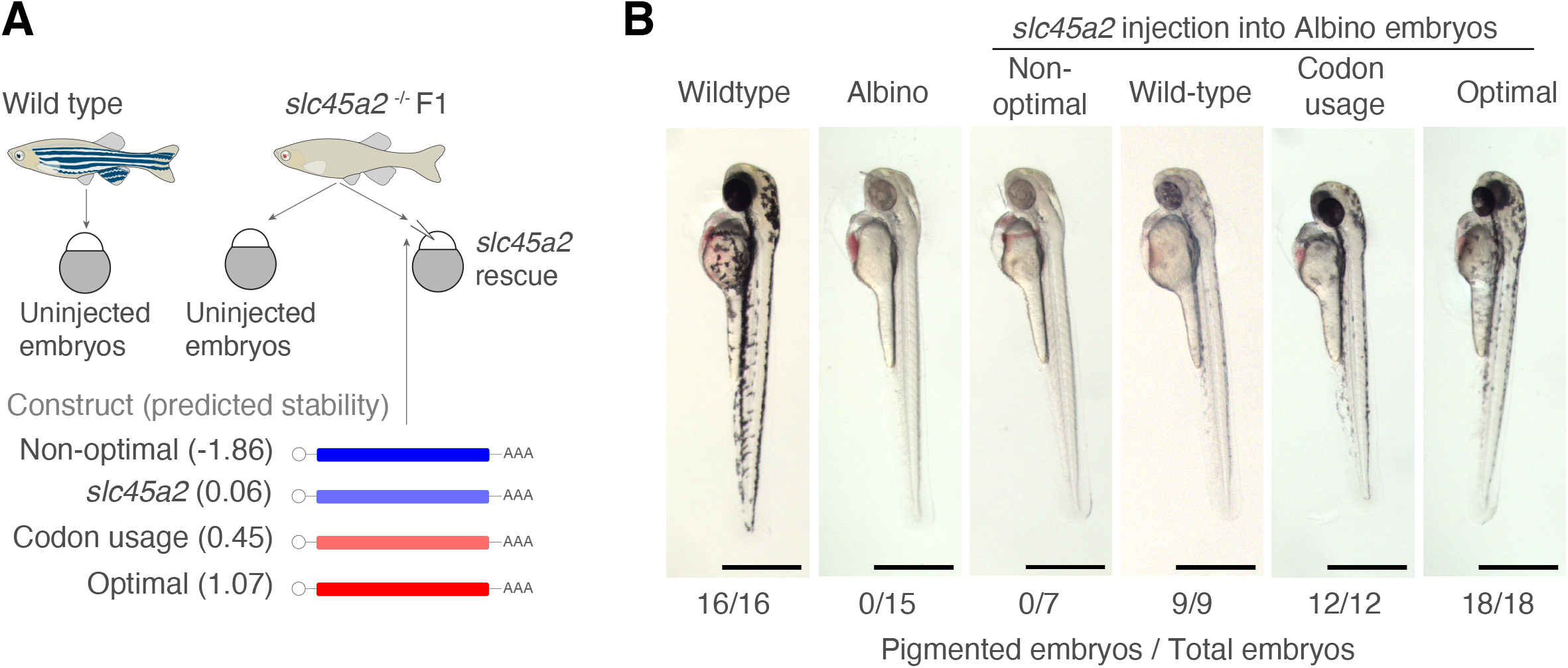
Optimized endogenous variant rescues loss-of-function phenotypes. **(A)** Schematic representation of the rescue experiment in zebrafish embryos. Variants of the *slc452* gene (melanin pigmentation) were injected into loss-of-function *slc45a2* knockout zebrafish embryos (albino phenotype, lack of pigmentation). The predicted mRNA stability of the variants is indicated. **(B)** Microscopy images of injected zebrafish embryos 48 hours post fertilization showing that only original and optimized variants rescued the loss-of-function phenotype. Scale bars represent 700 μm. The numbers reflect the proportion of embryos that showed melanin pigmentation.

### Steps to use iCodon

The iCodon interactive web interface is available at www.icodon.org as well as at https://bazzinilab.shinyapps.io/icodon/. The user can provide a coding sequence (A, T, C and G, case-insensitive) and select the relevant species (human, mouse, zebrafish or *Xenopus*) **(Figure 5A)**. A warning message is displayed if the pasted sequence is not a multiple of three, contains an internal stop codon, or does not contain a terminal stop codon. The results are displayed graphically, showing the degree of optimization achieved **(Figure 5B**), and the original sequence is plotted as a grey dot with the predicted stability. Nine optimized variants are shown in red and nine deoptimized sequences in blue, with their respective predicted stabilities indicated. Finally, by selecting the “download optimization results” link (**Figure 5C**), the user can retrieve the optimized and deoptimized coding sequences, as well as their predicted stabilities, number of codons and nucleotides changes in text format ready for downstream synthesis applications, (csv extension) (**Figure 5D**). Moreover, if the user wishes to study multiple sequences, iCodon can be run directly in R https://github.com/santiago1234/iCodon **(Supplemental file 1)**.

**Figure 5.**
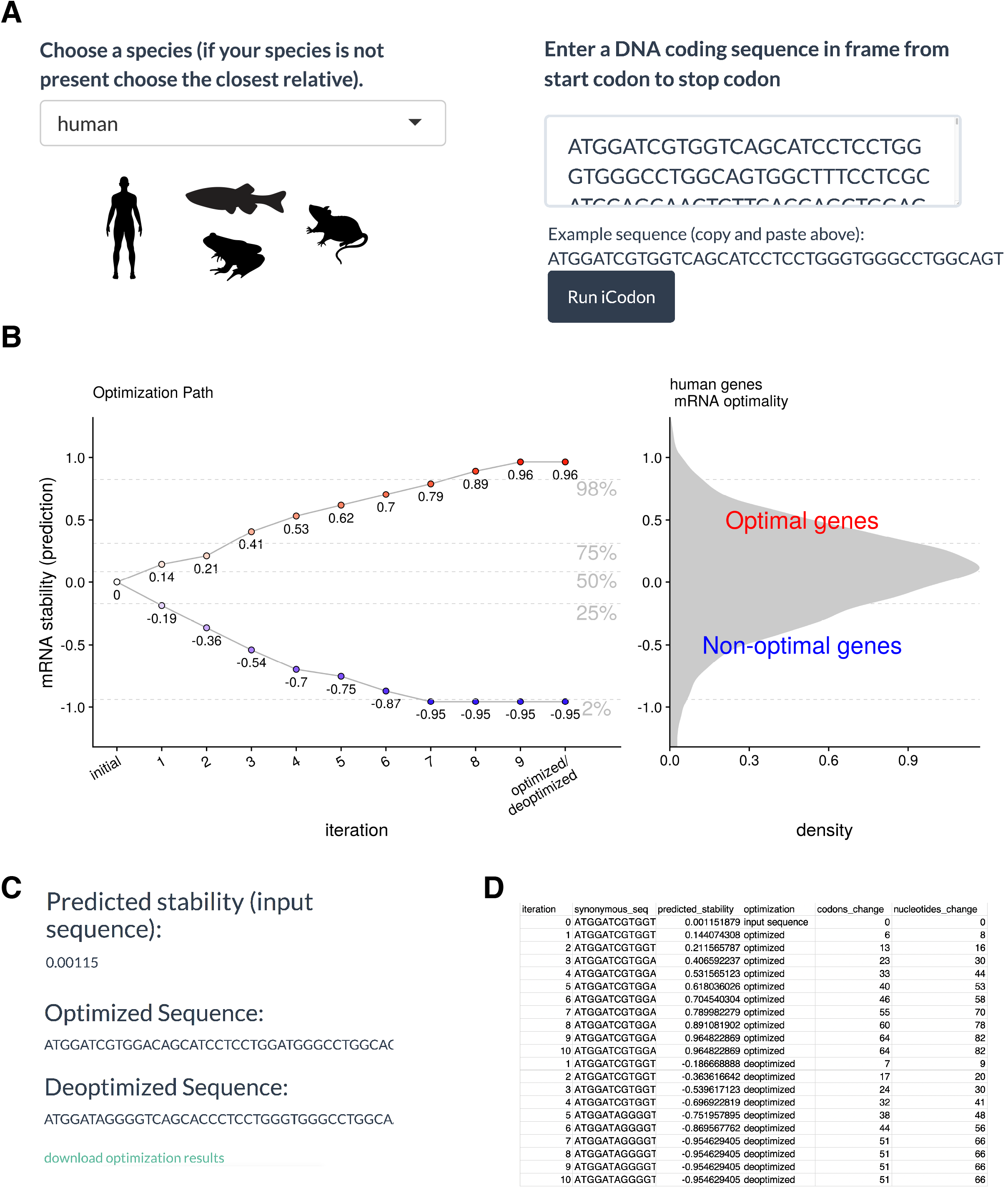
iCodon user steps. **(A)** The user needs to select between four species (human, mouse, zebrafish or *Xenopus*). The coding sequence (A, T, G and C) needs to be pasted into the box indicated and the Run iCodon box needs to be clicked. **(B)** The scatter plot will show the original sequence in grey with its predicted stability. Each of the optimized (red) or deoptimized (blue) sequences with each respective stability score will be displayed. A histogram of the mRNA stability distribution of endogenous genes of the selected species is shown to use as a reference for the designed variants. **(C)** The original sequence, as well as all designed variant sequences, stability scores and nucleotide/codon changes with respect to the original sequence will be provided in a file by clinking “download optimization results”. (**D**) Example table of the downloaded iCodon results.

## Discussion

The protein production outcome can be collectively influenced by regulatory elements encrypted in the promoter, 5′UTR, coding sequence, and 3′UTR (Nieuwkoop, Finger-Bou, van der Oost, & Claassens, 2020). Here, we have shown that iCodon can design *in silico* sequences with increased or decreased stability profiles by codon synonymous substitutions for vertebrates. We anticipate a number of applications for iCodon. First, and at the most basic level, users can interrogate the stability of endogenous genes based on their coding sequence. As mentioned above, knowing the relative stability of a gene of interest can provide potential insight into its biological role and/or evolution (e.g., inherently unstable mRNAs encoding core circadian components; **Figure 1C**) **(Figure 6)**.

**Figure 6.**
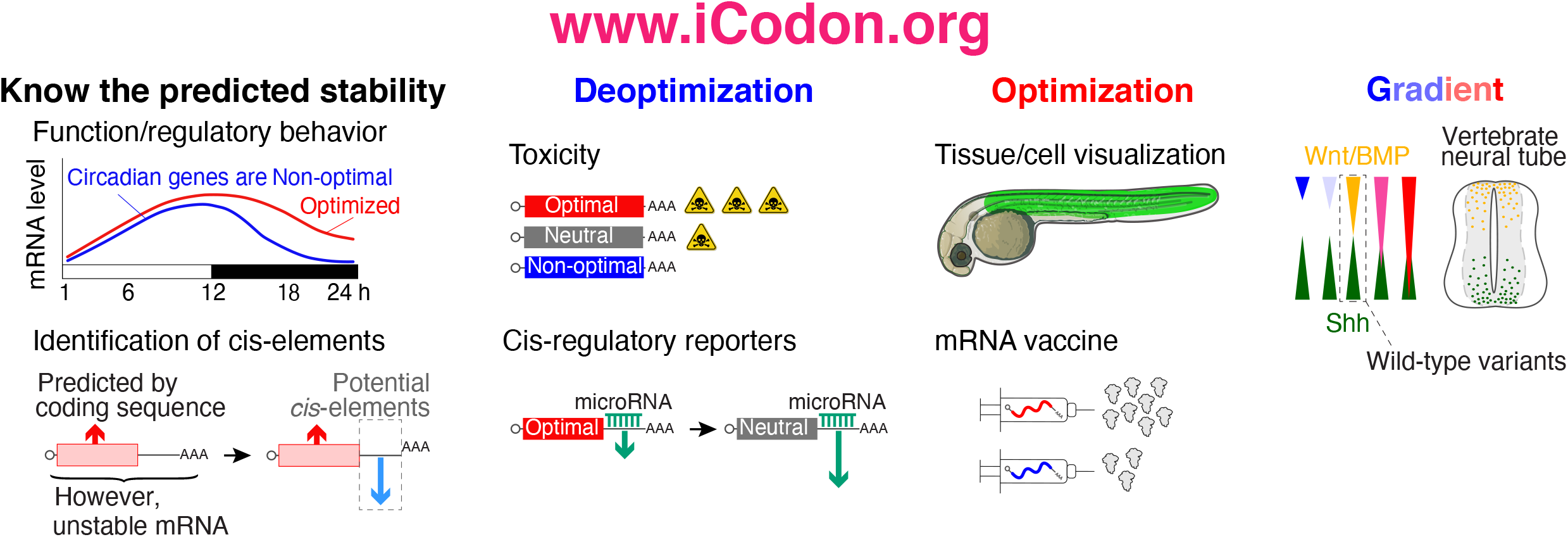
iCodon potential applications. iCodon can be used to uncover gene expression patters from the coding sequence; or to identify cis-regulatory elements. iCodon can be used to design sequences with neutral mRNA stability, these sequences will be more responsive to other regulatory elements (i.e., miR-430). Exogenous genes that are wanted to be expressed in a vertebrate model organism could result toxic for the cell; by designing coding sequences with a decreased expression, the toxicity level can be reduced. Tissue/cell visualization: fluorescent proteins found in another organism can be optimized for expression in vertebrate species. Injected mRNA (zebrafish embryos) or mRNA vaccine design can be codon optimized to increase the mRNA stability and expression. Expression gradient: iCodon has the potential to design a wide variety of coding sequences, which will show different gene expression patterns.

Second, we have recently shown that an accounting of codon-mediated effects on mRNA stability can help to identify other contributing *cis*-regulatory elements (Medina-Muñoz et al., 2021). For instance, microRNAs and RNA modifications such as m^6^A target a subset of maternal mRNAs for degradation during embryogenesis (Bazzini, Lee, & Giraldez, 2012; Bushati, Stark, Brennecke, & Cohen, 2008; Giraldez et al., 2006; Kontur, Jeong, Cifuentes, & Giraldez, 2020; Lund, Liu, Hartley, Sheets, & Dahlberg, 2009; Vastenhouw, Cao, & Lipshitz, 2019), whereas other modifications (e.g., m5C) are associated with stabilization of maternal RNAs (Y. Yang et al., 2019). Interestingly, we observed higher enrichment of destabilizing (miR-430 and m^6^A) or stabilizing (m^5^C) *cis*-regulatory elements in the 3′UTRs of genes in which codon optimality did not explain observed stability when compared to genes in which codon composition was highly predictive of observed stability (Medina-Muñoz et al., 2021). Hence, the predicted mRNA stability based on codon composition can be used to study other gene regulatory networks **(Figure 6)**.

Third, users can re-design the codon composition of reporter mRNAs and transgenes for specific downstream purposes. For example, we have observed that microRNA and m^6^A regulation dictate mRNA stability in conjunction with codon optimality (Medina-Muñoz et al., 2021). Specifically, microRNA or m^6^A targets with more optimal coding sequences are more stable than non-optimal target mRNAs (Medina-Muñoz et al., 2021). However, microRNAs (miR-430/-427) targeting efficacy is reduced in genes highly enriched in optimal or non-optimal codons during embryogenesis (Medina-Muñoz et al., 2021). Therefore, researchers might want to evaluate the codon optimality of classical GFP, mCherry or luciferase reporters and subsequently deoptimize them to reach an average level of stability (Q. Wu et al., 2019) **(Figure 6)**. Such re-tooling may be well-advised, as some ‘highly-stable’ reporter mRNAs may not reflect the majority of endogenous stability profiles. Additionally, reducing the level of gene expression can be desired to reduce the toxic effect of highly expressed proteins (**Figure 6**).

Fourth, users can design transgenes to be more highly expressed for a myriad of applications, including protein visualization (e.g., GFP or AausFP1, **Figure 2-3**), mRNA knock-down (Kushawah et al., 2020), genome editing (Moreno-Mateos et al., 2017) and loss-of-function rescue (**Figure 4 and 6)**. Through iCodon optimization, the amount of mRNA required to rescue loss-of-function phenotypes can be reduced, thereby avoiding toxic effects associated with the introduction of high amounts of exogenous RNA (Tsetskhladze et al., 2012). While screening random synonymous substitution libraries for variants with desired expression levels is a valid approach (Kudla, Murray, Tollervey, & Plotkin, 2009), the number of potential variants can be astronomical, depending on the coding sequence size. Therefore, iCodon provides a practical first step toward more targeted solutions.

Fifth, users may employ iCodon to design a range of variant transcripts that have slightly different expression profiles in order to, for example, measure dosage effects of morphogens during development (e.g., Sonic hedgehog (Shh), BMP or Wnt) **(Figure 6)**. Additionally, iCodon can simply be used when genes from one species need to be expressed in a heterologous context (e.g., AausFP1 from *Aequorea. cf. australis* into human cells and zebrafish embryos, **Figure 3**).

Finally, we envision the use of iCodon in the design of RNA-based therapeutics (e.g., mRNA vaccines) (**Figure 6)**, in which increased stability and expression may correlate with stronger efficacy (e.g., stronger immune response) and/or smaller doses (Jackson, Kester, Casimiro, Gurunathan, & DeRosa, 2020; Krienke et al., 2021; Pardi, Hogan, Porter, & Weissman, 2018).

While iCodon predictions correlated with observed gene expression for most genes tested, we have observed that particular variants have not necessarily followed the expected expression level. As stated above, there is other regulatory information encrypted in the coding sequence that can affect mRNA stability and gene expression, such as translational ramp (Verma et al., 2019), lysine homopolymers (Koutmou et al., 2015), and/or protein folding/activity (Yu et al., 2015), just to mention a few. Therefore, we recommend designing more than one synonymous sequence, as optimization requires experimental validation.

## Conclusion

In summary, iCodon provides a simple tool for the scientific community to interrogate mRNA stability of their genes of interest based on codon composition, and to design strategies to modulate expression levels in vertebrates through codon optimization or deoptimization.

## Abbreviations

mRNA: messenger RNA
miR: microRNAs
m^6^A: N6-methyladenosine
m^5^C: 5-methylcytosine
P2A: 2A ribosome skipping sequence
GFP: green fluorescent protein
EGFP: enhanced green fluorescent protein
AausFP1: *Aequorea. cf. australis* fluorescent protein 1
UTR: untranslated regions

## Declarations

### Ethics approval and consent to participate

Does not apply

### Consent for publication

The authors declare no competing interests.

### Availability of data and materials

The source code and datasets generated during the current study are available in the GitHub repository, https://github.com/santiago1234/iCodon. The iCodon interactive web interface is available at www.iCodon.org or https://bazzinilab.shinyapps.io/icodon/.

### Authors’ contributions

AAB and SGMM conceived the study. SGMM developed and implemented the iCodon algorithm. MD and LAC conceived, planned and performed the experimental validation. GdSP performed all the zebrafish experiments. QW performed experiments in human cells. AAB supervised the work. SGMM developed the iCodon web interface with input from the other authors. SGMM, MD, LAC and AAB wrote the manuscript with input from the other authors. The authors read and approved the final manuscript. This work was performed as part of thesis research for LAC, GdSP, and QW, Graduate School of the Stowers Institute for Medical Research.

## Acknowledgments

We thank Dr Carter Takacs, Dr Cei Abreu-Goodger, and Dr Daniel Cifuentes for suggestions and critical reading of the manuscript. We also thank all Bazzini lab members for their help, and the following core facilities for their support: Molecular Biology, Aquatics, Cytometry, Microscopy, Tissue Culture, and Media Prep.

---

**Supplemental figure 1.**
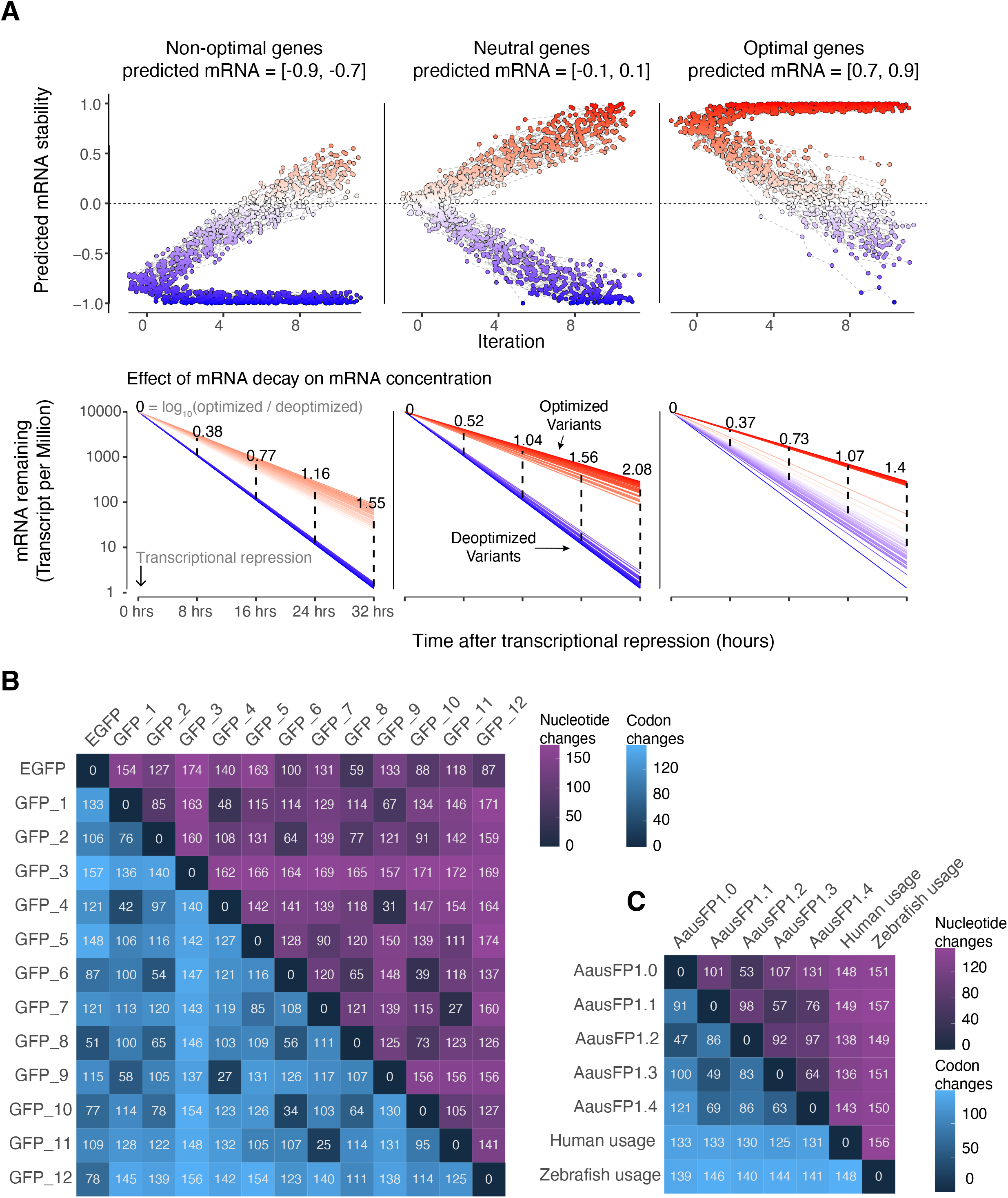
(**A**) Optimization of endogenous genes. Three sets of genes were selected (50 genes in each group): genes that are predicted to be non-optimal (left), genes predicted to be neutral (middle), and genes predicted to be optimal (right). For each of these genes, we ran iCodon to optimize and deoptimize them. The top row shows the iCodon optimization results. The x-axis is the iteration number, and the y-axis is the predicted mRNA stability. The circles connected by a dashed-line show the optimization or deoptimization path for each gene. Gaussian noise was added to the points. The bottom row shows a simulation of the effect of mRNA stability optimization achieved on mRNA abundance. At time 0 h transcription is inhibited, and the genes decay according their predicted stability. The lines in red color denote the optimized genes, the lines in blue are the deoptimized genes. Messenger RNA abundance (Transcript per Million) is plotted on a logarithmic scale. The points and dashed lines show the predicted change in expression (log10 fold change) between optimized and deoptimized variants. (**B**) Matrix comparing the pairwise distance between the synonymous GFP variants. The lower diagonal entries show the distance in codon changes and the upper diagonal entries in nucleotide changes. (**C**) Same as in panel B for AausFP1variants.

## Methods

### Evolutionary algorithm for codon optimization

The number of possible synonymous sequences coding for the same peptide is astronomically large. To solve the problem of finding a particular sequence with a target mRNA stability, we developed a genetic algorithm [1]. This algorithm operates on three steps (**Figure 1c**):

**Initialization Step:** An initial coding DNA sequence is provided together with a vertebrate species (human, mouse, zebrafish, or *Xenopus*). Also, a fixed threshold ***t*** is set (***t*** = 1 by default). This threshold represents the maximum predicted mRNA stability that a sequence can achieve.

**Evolution Step**: ***N*** daughter sequences are generated (***N*** = 10 by default). For each random sequence, a proportion of ***p*** codon positions (***p*** = 0.05 by default) are selected uniformly at random (excluding start and stop codons). For each of these positions, a synonymous codon is randomly selected to introduce a synonymous mutation. We created a custom sampling distribution for selecting random synonymous codons. For the optimization process the distribution samples optimal codons with higher frequency and for the deoptimization process non-optimal codons are sampled with higher frequency. This sampling distribution was generated by ranking the codon stabilization scores [2–4] and then applying the softmax function.

**Selection Step**: The fitness of each daughter sequence is evaluated, and the fittest sequence is selected based on whether the daughter follows the optimized or deoptimized path. The fitness of the sequence is the predicted mRNA stability based on the codon composition [5].

This process is iterated ***m*** times (***m*** = 10 by default) producing an optimization path. In each iteration the fittest sequences are kept (i.e., the most optimal and the most non-optimal). The optimization path produces a gradient in mRNA stability level. If the predicted mRNA stability of the fittest daughter sequence is more than the threshold ***t***, the process will stop generating new daughter sequences. The last sequences in the iteration correspond to the most optimized or deoptimized variants. The default parameters were selected by heuristic observations.

iCodon is implemented in R (version >= 3.6.2), and the source code is available from GitHub https://github.com/santiago1234/iCodon.

### Gene Ontology term analysis

The Gene Ontology term analysis was conducted on the GOrilla website http://cbl-gorilla.cs.technion.ac.il/ [6], with two unranked lists of genes as running mode. Either the Top 100 optimal or non-optimal genes were selected as target, and the rest of the genes were selected as background.

### Transcriptome analysis

The coding genes in each transcriptome were downloaded from biomart (Ensembl Genes 102, human = GRCh38.p13, *Xenopus* = Xenopus_tropicalis_v9.1, mouse = GRCm38.p6, and zebrafish = GRCz11) [7]. The longest isoform for each gene was kept. The predicted mRNA stability for each gene was computed with iCodon [5] (Supplemental tables 1-4).

For the stability comparison between groups of genes in human (Figure 1E), circadian genes were chosen based on the core components of the circadian clock [8], histones and zinc finger proteins were obtained from Uniprot under the search terms ‘histone’ and ‘zinc finger protein’, and curated manually, and ribosomal protein genes were obtained from the Gene Ontology term structural constituent of ribosome (GO:0003735).

### Variants clones

Sequences designed by either iCodon or IDT Codon Optimization Tool were synthetized by IDT and cloned into pCS2 backbones using conventional restriction cloning or HiFi DNA Assembly (NEB). Synonymous reporters (Figure 1 A-B) were previously constructed [4] with the exception of the Codon Usage synonymous reporter that was cloned as described above. All sequences can be accessed in Supplemental Table 6.

### Human cells transfection experiments

293T cells were obtained from the Tissue Culture core facility at the Stowers Institute for Medical Research. For transfection, cells were plated in 96-well plates at a relatively low passage, cultured with DMEM media, 10% FBS, L-glutamine and penicillin/streptomycin, and set overnight to reach 70% confluency the day of transfection. Prior transfection, all plasmids were quantified using the Qubit Fluorometric Quantification. 293T cells were transfected using Lipofectamine 3000 based on the manufacturer’s instructions, and 24 hours post transfection, cells were collected for cytometry analysis. The fluorescence intensity of the cells was quantified in a ZE5 Cell Analyzer, using lasers and detectors for GFP (488/510) and mCherry (587/610). The cytometry data .fsc file were analyzed with FCS Express 7, and the median intensity of the cells was used to represent fluorescence intensity.

### mRNA in vitro transcription

Plasmids carrying the different constructs employed in this study were first digested to linearize the DNA and then used for in vitro transcription using the mMESSAGE mMACHINE® Kit, following manufacturer’s protocols. Prior injections, mRNA was quantified using the Qubit Fluorometric Quantification.

### Zebrafish Embryo Injection and Image Acquisition

Optimized and deoptimized mRNAs encoding EGFP (100pg), AausFP1 (100pg), or *slc45a2* (200pg) were co-microinjected with TagRFP (100pg) in zebrafish embryos at 1-cell stage (see figure legends for details in each experiment). Injected embryos were collected, mounted for imaging in low melting point agarose as described in [9], analyzed and quantified between 24 hours and 2 days post fertilization depending on the experiment.

Zebrafish embryo fluorescent pictures were analyzed using a Nikon TI2-E inverted microscope, photographed with a Photometrics Prime 95B-25MM back-illuminated sCMOS camera using laser for GFP (491/535) and RFP (574/615) and images processed with NIS-Elements Advanced Research Package software. Images were further processed in Fiji. AausFP1, GFP and RFP fluorescence were quantified using Fiji (Image J) software by performing sum slices z-projection, selecting the embryo’s trunk region and measuring mean pixel value in the same manner for all conditions.

Zebrafish embryo pigmentation phenotype pictures were analyzed using a ‘Zeiss Lumar V12 steREO’ microscope with same conditions for all the injected embryos and images processed with MicroManager software version 1.4.23 [10].

### Albino zebrafish line

To generate the albino loss-of-function zebrafish, 1-cell stage embryos were injected with Cas9 mRNA along with gRNA targeting the coding sequence of *slc45a2* as previously reported [11]. The F0 lacked melanin pigmentation and were crossed to obtain the F1 used for the experiments of this study.

### Zebrafish Maintenance

Zebrafish experiments were done according to the IACUC approved guidelines. Zebrafish embryos collected for microinjections were coming from random parents (AB, TF and TLF, 6-25 months old) mating from 3 independent strains or from random parents (*slc45a2^−/−^* in AB, TF and TLF background, 6-25 months old) mating from 1 strain. The embryos were pooled from random 12 males and 12 females for each set of experiments, or random 8 males and 3 females from the *slc45a2^−/−^* parents.

### Statistical analysis

All statical analyses were conducted in R version 4.0.2. Displayed data results from at least two independent experiments.

